# DrosOmics: a comparative genomics browser to explore omics data in natural populations of *D. melanogaster*

**DOI:** 10.1101/2022.07.22.501088

**Authors:** Marta Coronado-Zamora, Judit Salces-Ortiz, Josefa González

**Affiliations:** Institute of Evolutionary Biology, CSIC, Universitat Pompeu Fabra, Barcelona, Spain

## Abstract

The advent of long-read sequencing technologies has allowed the generation of multiple high-quality de novo genome assemblies for multiple species, including well-known model species such as *Drosophila melanogaster*. Genome assemblies for multiple individuals of the same species are key to discover the genetic diversity present in natural populations, especially the one generated by transposable elements, the most common type of structural variant. Despite the availability of multiple genomic datasets for *D. melanogaster* populations, we lack an efficient visual tool to display different genomes assemblies simultaneously. In this work, we present DrosOmics, a comparative genomics-oriented browser for 52 high-quality reference genomes of *D. melanogaster*, including annotations from a highly reliable set of transposable elements, and functional transcriptomics and epigenomics data for half the populations. DrosOmics is based on JBrowse 2, which allows the visualization of multiple assemblies at once, key to unraveling structural and functional features of *D. melanogaster* natural populations.

## Introduction

The advent of technological developments in DNA sequencing have led to a high number of whole-genome sequences for multiple individuals of the same species, revolutionizing the field of comparative population genomics (Mitsuhashi and Matsumoto 2020; Sakamoto et al. 2020). Among these developments, long-read sequencing technologies allowed to improve the quality and completeness of reference genomes because of their ability to span repetitive regions, leading to a better detection and annotation of structural variants such as transposable elements (TEs) (Solares et al. 2018; Du and Liang 2019; Miga et al. 2020; Rech et al. 2022). *Drosophila melanogaster*, one of the central model species for molecular population genomics studies (Casillas and Barbadilla 2017) and for studying TEs (Mccullers and Steiniger 2017), have benefited from these technological developments and currently has several *de novo* reference genomes representative of natural populations sampled worldwide (Long et al. 2018; Chakraborty et al. 2019; Rech et al. 2022).

Complementary to these population genome sequence data, there has been a large increase in the generation of other functional -omics data, such as transcriptomics and epigenomics. While the genome sequence represents a static component of the organism, functional -omics data vary across different body parts or tissues (spatial), life stages (temporal), or conditions (*e.g*. different environments and experimental treatments). Capturing this information is relevant for *e.g*. understanding how organisms adapt to changing environments, plant and animal breeding, and medical genetics.

One major challenge for genomic studies is to visualize and retrieve large amounts of genome and functional -omics data. For that, web-based genome browsers are a suitable platform that allows browsing, visualizing and retrieving data through an efficient and user-friendly graphical interface (Schattner 2008; Buels et al. 2016). Current population genomics-oriented browsers are based on the first available JBrowse version, which only allows a single genome to be displayed. These browsers, *e.g*. PopHuman (https://pophuman.uab.cat; Casillas et al. 2018), were designed for visualizing and querying population genetic statistics for each of the populations included. For *D. melanogaster*, there are two such population genomics-oriented browsers available: PopFly (https://popfly.uab.cat; Hervas et al. 2017), and DEST (https://dest.bio; Kapun et al. 2021). However, until recently, JBrowse-based browsers preclude the comparative analysis of sequence and structural variants across genomes.

## New approaches

In this work, we present DrosOmics, a comparative genomics-oriented browser based on the new version of JBrowse, JBrowse 2 (Diesh et al. 2022), that allows browsing, visualizing and retrieving genome sequences and functional -omics data, for 52 genomes of *D. melanogaster* collected in 32 locations worldwide (Table 1). JBrowse 2 allows to explore the sequence and structural variation of several populations simultaneously and to quickly navigate through the different genome views. The genomes included in DrosOmics have been annotated with a high-quality and manually-curated library of transposable elements (TEs), which includes the sequence variability present in several natural populations (Rech et al. 2022). DrosOmics also compiles functional annotations for half of the genomes, including transcriptomics (RNA-seq) and epigenomics datasets (ChIP-seq and ATAC-seq). This allows genetic variants, *e.g*., single nucleotide polymorphisms (SNPs) and structural variants, such as TEs, to be associated with variation in gene expression and epigenetic modifications.

## DrosOmics browser content overview

DrosOmics compiles 52 high-quality genome assemblies that have been obtained using long-read sequencing technologies (Table 1 and Table S1).

**Table 1.** 52 *D. melanogaster* reference genome assemblies compiled in DrosOmics.

| Genomes | Source locations | Data source |
| --- | --- | --- |
| 32 genomes | 12 localities from 8 countries: America (1), Europe (7) | Rech et al. (2022) |
| 14 genomes | 14 localities from 11 countries: Africa (2), Europe (2), North America (1), North Atlantic Ocean (1), South America (2), Asia (3) | Chakraborty et al. (2019) |
| 5 genomes | 5 localities from 5 countries: Africa (1), Europe (1), America (1), Asia (1), Oceania (1) | Long et al. (2018) |
| 1 genome (ISO-1) | America | Solares et al. (2018) |

DrosOmics contains gene and transposable elements (TEs) annotations for the aforementioned 52 *D. melanogaster* reference genomes. Note that the current version of DrosOmics provides gene annotations that were transferred to each genome from the reference release 6 of *D. melanogaster*. Functional -omics data for 26 genomes were retrieved from previous research (Everett et al. 2020; Salces-Ortiz et al. 2020; Green et al. 2022; Horváth et al. 2022; Table 2 and Table S1) or are reported in this work for the first time (Table 2, Table S1 and Supplementary Material). Genome and functional annotations can be graphically displayed along the chromosome arms for multiple genomes simultaneously (Table 2). DrosOmics is built on JBrowse 2 (release 1.6.9) (Diesh et al. 2022) and is currently running under a docker container using Apache on a CentOS 7.9.2009 Linux x64 server with 4 vCores processors and 16 GB RAM.

**Table 2.** Available tracks in DrosOmics: >500 available tracks. Further details on raw data number accessions, data processing and publications associated are available on Supplementary Material and Table S1. Symbols indicate the publication source for the functional data: ◆ This publication; ■ Everett et al. (2020); ▲ Salces-Ortiz et al. (2020); ● Green et al. (2022); ► Horváth et al. (2022). The number of biological replicates (repl) for each dataset is given. FC: fold change.

| Track type | Track name | Description | Genomes | Datasets |
| --- | --- | --- | --- | --- |
| Genome sequence | Reference sequence | Displayed by chromosome arm (2R, 2L, 3L, 3R and X) | All | 52 |
| Genome annotations | TE annotations<br>Gene annotations | - *De novo* TE annotations<br>- Gene annotations transferred from *D. melanogaster* release 6 annotation | All | 104 |
| RNA-seq | RNA-seq coverage | - Basal conditions: head, gut, ovary<br>- Basal conditions: whole-body<br>- Basal and malathion treatment: gut<br>- Basal and copper treatment: whole body ♀<br>- Basal and desiccation treatment: whole body ♀ | ♦ 5 genomes<br>■ 8 genomes<br>▲ 4 genomes<br>● 6 genomes<br>► 6 genomes | 15 (3 repl)<br>8 (2 repl)<br>8 (3 repl)<br>12 (3 repl)<br>12 (3 repl) |
| ChIP-seq | ChIP-seq FC signal<br>ChIP-seq peak calling | - H3K9me3 (repressive mark): head, gut, ovary<br>- H3K27ac (active mark): head, gut, ovary | ♦ 5 genomes<br>♦ 5 genomes | 15 (3 repl)<br>15 (3 repl) |
| ATAC-seq | ATAC-seq FC signal<br>ATAC-seq peak calling | - Basal and malathion treatment whole-body ♀ | ▲ 4 genomes | 8 (3 repl) |

DrosOmics has a Help section including documentation about all the data compiled in the browser, such as detailed data descriptions, data sources and citations, as well as details about the browser tracks available. Importantly, it also provides an exhaustive Tutorial with step-by-step worked examples introducing the usage of the browser. All the compiled data and support resources provided by DrosOmics are open and available at http://gonzalezlab.eu/drosomics.

## DrosOmics case studies

To show the types of questions and analyses that DrosOmics facilitates, we have selected three case studies to demonstrate the usability of the browser: i) *analysis of a region of interest in multiple genomes;* ii) *analysis of structural variability between genomes*; and iii) *study of the association between structural, epigenetic and expression data*.

### Case study 1

With DrosOmics, the user is able to analyze and retrieve the sequence of a region of interest in any of the 52 genomes available. This is of special interest for the study of the presence and/or absence of genetic variants in natural populations. One example is the analysis of the presence of a 49-bp indel polymorphism, and a linked SNP, in the 3’UTR region of the *metallothionein A* (*MtnA*) gene in some natural populations. This indel has been associated with higher expression of *MtnA*, linked, in turn, to oxidative stress tolerance (Catalán et al. 2016). With DrosOmics, the user can navigate to the *MtnA* gene region and visualize the sequence of the 3’UTR from any of the genomes to determine if they carry the indel or not (Figure 1). This case study exemplifies the utility of the browser in visualizing a sequence from multiple genomes for validation and further characterization.

**Figure 1.**
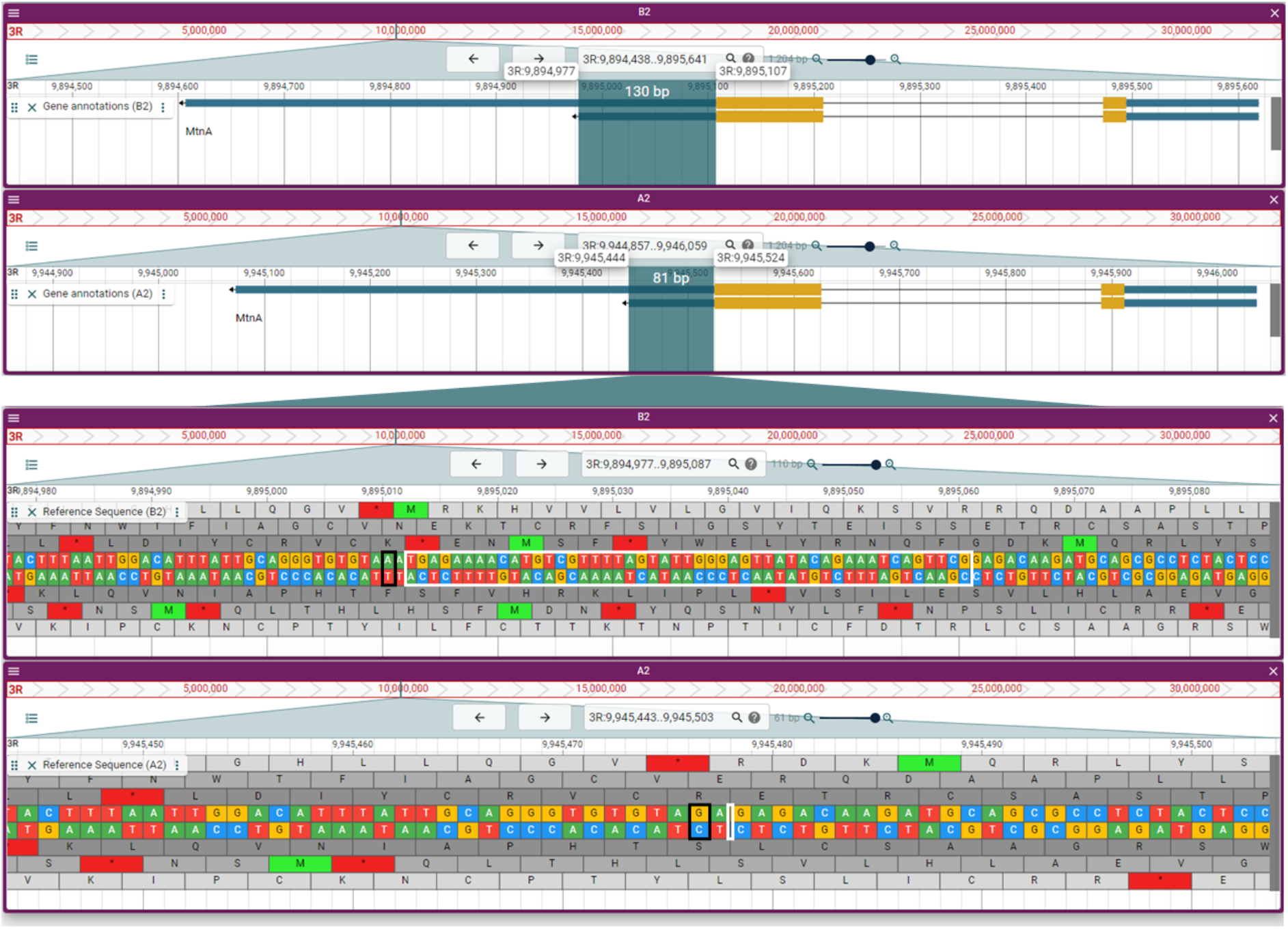
DrosOmics snapshot showing *mtnA* gene and its 3’UTR sequence in two genomes. Genome view of the *MtnA* gene from genome *B2* (top; Cape Town (South Africa); Chakraborty et al. 2019) and genome *A2* (bottom; Bogota (Colombia); Chakraborty et al. 2019). By zooming to the sequence-level in the 3’UTR of *mtnA* gene, it is possible to visualize the 49-bp indel (highlighted in white) present in *B2* and absent in *A2*, and the linked SNP (highlighted in black) described in Catalán et al. (2016).

### Case study 2

One of the main advantages of JBrowse 2 is that it allows the study of the structural variation between genomes. To exemplify this, we will focus on the genome region surrounding the *Cyp6a17* gene, which was found to be associated with increased cold preference when knockdown (Kang et al. 2011). Chakraborty et al. (2019) described that *Cyp6a17* was not present in a high proportion of African and North American flies, suggesting an adaptive role of this mutation on temperature regulation in the wild. With DrosOmics it is possible to identify and visualize genomes with the present and absent genotypes. Figure 2 shows four genomes from North American *D. melanogaster*: two (RAL-176 and RAL-375) have the copy of *Cyp6a17* and the other two (RAL-059 and RAL-177) do not. By including RNA-seq coverage tracks, it is possible to visualize the expression differences of the genes in the two genotypes: when present, *Cyp6a17* and their neighboring paralogs –*Cyp6a22* and *Cyp6a23*–exhibit similar expression levels. When *Cyp6a17* is absent, the expression level of *Cyp6a23* is notably higher than that of *Cyp6a22*. With this example, we show that DrosOmics can be a tool to easily explore the structural differences between genomes and their possible functional consequences.

**Figure 2.**
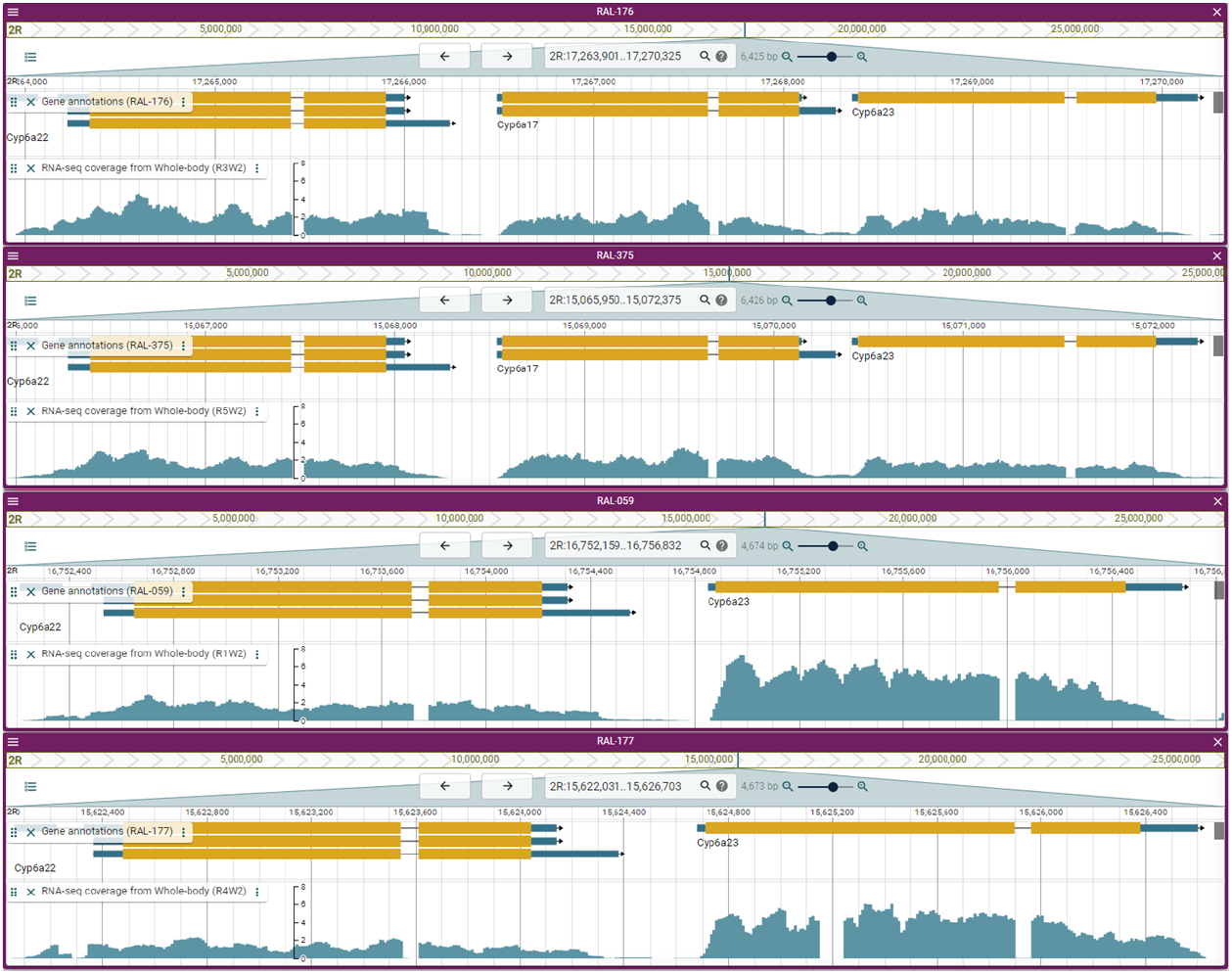
DrosOmics snapshot showing the region surrounding *Cyp6a17* gene in four genomes from North American *D. melanogaster*. RAL-176 and RAL-375 (top) have *Cyp6a17* and RAL-059 and RAL-177 (bottom) have the absent genotype. RNA-seq profiles are shown in turquoise.

### Case study 3

This example links gene expression changes with the epigenetic effects of a transposable element. TEs have been associated with gene downregulation due to the spread of repressive histone marks targeting TEs (Sienski et al. 2012; Choi and Lee 2020). Figure 3 shows the region surrounding the gene *Myo31DF* in two genomes: TOM-007 and JUT-011 (Rech et al. 2022). TOM-007 has the insertion of a TE that is not present in JUT-011 genome. By activating ChIP-seq tracks for the repressive histone mark H3K9me3 (fold enrichment [FC] and histone peaks) one can observe the enrichment and presence of peaks surrounding the TE insertion that are not present when the TE is absent. Because the TE is in the promoter region of *Myo31DF*, this epigenetic effect can have an impact on the gene expression level. Indeed, the RNA-seq levels of the gene with the insertion are lower than when the insertion is absent (*z*-score = −25.4, *p*-value < 0.001; see Supplementary Materials; Coronado-Zamora and González, *in prep*.).

**Figure 3.**
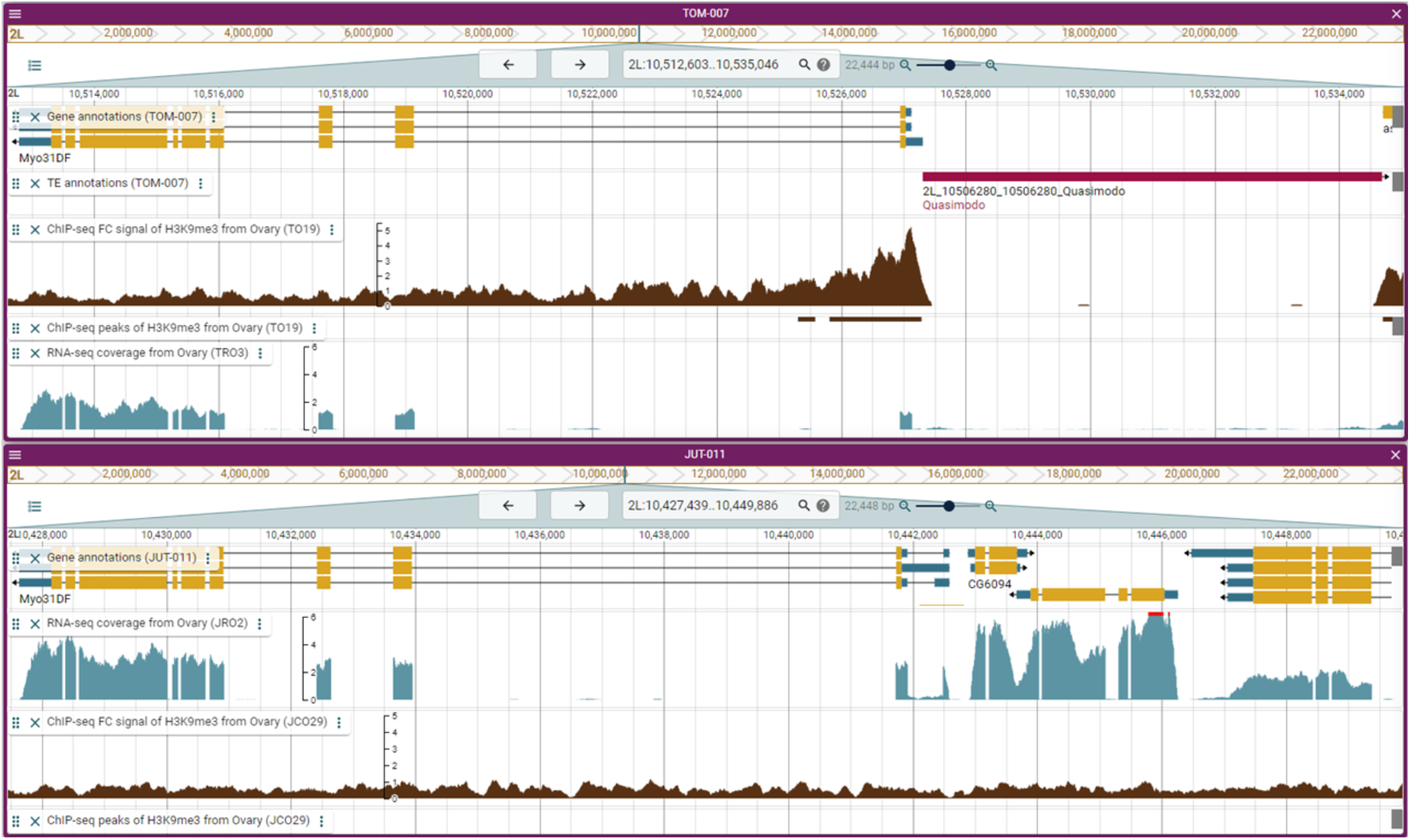
DrosOmics snapshot showing the region surrounding *Myo31DF* gene in two genomes: TOM-007 (with a TE insertion in the promoter region) and JUT-11 (no TE insertion). The presence of the TE induces the enrichment of H3K9me3, affecting the promoter region of *Myo31DF*. This leads to the downregulation of the gene.

## Conclusions

The DrosOmics browser, based on JBrowse 2, compiles genome sequences, transposable elements and gene annotations, as well as functional -omics data for 52 genomes from *D. melanogaster* natural populations. Furthermore, as JBrowse 2 is a highly scalable platform, it allows the continuous update by the addition of new annotations, populations, or new *Drosophila* species. We believe that DrosOmics will facilitate the unraveling of structural and functional features of *D. melanogaster* natural populations, thus providing a great opportunity to explore and test new evolutionary biology hypotheses. DrosOmics is open and freely available at http://gonzalezlab.eu/drosomics.

## Data availability

All processed data is available to download from http://gonzalezlab.eu/drosomics. The RNA-seq and ChIP-seq raw data for five genomes that are reported here for the first time are available in the NCBI Sequence Read Archive (SRA) database under BioProject PRJNA643665.

## Acknowledgements

We would like to thank the UPF Scientific IT Core Facility for support on implementing the JBrowse platform, Santiago Radío (from González Lab) for providing the TE and gene annotations for the five genomes of Long et al. (2018), and the DrosEU consortium for sharing flies with us.

## Funding

This work was supported by the European Research Council (ERC) under the European Union’s Horizon 2020 research and innovation programme (H2020-ERC-2014-CoG-647900), and by grant PID2020-115874GB-I00 funded by MCIN/AEI/10.13039/501100011033. DrosEU is funded by a Special Topic Network award from the European Society for Evolutionary Biology (ESEB). J.S.-O. was funded by a Juan de la Cierva-Formación fellowship (FJCI-2016-28380).

## Notes

http://gonzalezlab.eu/drosomics

### Competing Interest Statement

The authors have declared no competing interest.

### Summary of Updates

We have added case studies to show the types of questions and analyses that the comparative genomics browser DrosOmics facilitates.

## References

Buels R, Yao E, Diesh CM, Hayes RD, Munoz-Torres M, Helt G, Goodstein DM, Elsik CG, Lewis SE, Stein L, et al. 2016. JBrowse: A dynamic web platform for genome visualization and analysis. Genome Biol. 17:1–12.

Casillas S, Barbadilla A. 2017. Molecular Population Genetics. Genetics 205:1003–1035.

Casillas S, Mulet R, Villegas-Mirón P, Hervas S, Sanz E, Velasco D, Bertranpetit J, Laayouni H, Barbadilla A. 2018. PopHuman: the human population genomics browser. Nucleic Acids Res. 46:D1003–D1010.

Catalán A, Glaser-Schmitt A, Argyridou E, Duchen P, Parsch J. 2016. An Indel Polymorphism in the MtnA 3’ Untranslated Region Is Associated with Gene Expression Variation and Local Adaptation in Drosophila melanogaster. PLOS Genet. 12:e1005987.

Chakraborty M, Emerson JJ, Macdonald SJ, Long AD. 2019. Structural variants exhibit widespread allelic heterogeneity and shape variation in complex traits. Nat. Commun. 2019 101 10:1–11.

Choi JY, Lee YCG. 2020. Double-edged sword: The evolutionary consequences of the epigenetic silencing of transposable elements. PLOS Genet. 16:e1008872.

Diesh C, Stevens GJ, Xie P, Martinez TDJ, Hershberg EA, Leung A, Guo E, Dider S, Zhang J, Bridge C, et al. 2022. JBrowse 2: A modular genome browser with views of synteny and structural variation. bioRxiv:2022.07.28.501447.

Du H, Liang C. 2019. Assembly of chromosome-scale contigs by efficiently resolving repetitive sequences with long reads. Nat. Commun. 2019 101 10:1–10.

Everett LJ, Huang W, Zhou S, Carbone MA, Lyman RF, Arya GH, Geisz MS, Ma J, Morgante F, St Armour G, et al. 2020. Gene expression networks in the Drosophila Genetic Reference Panel. Genome Res. 30:485–496.

Green L, Coronado-Zamora M, Radio S, Rech GE, Salces-Ortiz J, González J. 2022. The genomic basis of copper tolerance in Drosophila is shaped by a complex interplay of regulatory and environmental factors. bioRxiv:2021.07.12.452058.

Hervas S, Sanz E, Casillas S, Pool JE, Barbadilla A. 2017. PopFly: the Drosophila population genomics browser. Bioinformatics 33:2779–2780.

Horváth V, Guirao-Rico S, Salces-Ortiz J, Rech GE, Green L, Aprea E, Rodeghiero M, Anfora G, González J. 2022. Basal and stress-induced expression changes consistent with water loss reduction explain desiccation tolerance of natural Drosophila melanogaster populations. bioRxiv:2022.03.21.485105.

Kang J, Kim J, Choi K-W. 2011. Novel cytochrome P450, cyp6a17, is required for temperature preference behavior in Drosophila. PloS One 6:e29800.

Kapun M, Nunez JCB, Bogaerts-Márquez M, Murga-Moreno J, Paris M, Outten J, Coronado-Zamora M, Tern C, Rota-Stabelli O, Guerreiro MPG, et al. 2021. Drosophila Evolution over Space and Time (DEST): A New Population Genomics Resource. Mol. Biol. Evol. 38:5782–5805.

Long E, Evans C, Chaston J, Udall JA. 2018. Genomic Structural Variations Within Five Continental Populations of Drosophila melanogaster. G3 GenesGenomesGenetics 8:3247–3253.

Mccullers TJ, Steiniger M. 2017. Transposable elements in Drosophila. Mob. Genet. Elem. 7:1–18.

Miga KH, Koren S, Rhie A, Vollger MR, Gershman A, Bzikadze A, Brooks S, Howe E, Porubsky D, Logsdon GA, et al. 2020. Telomere-to-telomere assembly of a complete human X chromosome. Nat. 2020 5857823 585:79–84.

Mitsuhashi S, Matsumoto N. 2020. Long-read sequencing for rare human genetic diseases. J. Hum. Genet. 65:11–19.

Rech GE, Radío S, Guirao-Rico S, Aguilera L, Horvath V, Green L, Lindstadt H, Jamilloux V, Quesneville H, González J. 2022. Population-scale long-read sequencing uncovers transposable elements associated with gene expression variation and adaptive signatures in Drosophila. Nat. Commun. 2022 131 13:1–16.

Sakamoto Y, Sereewattanawoot S, Suzuki A. 2020. A new era of long-read sequencing for cancer genomics. J. Hum. Genet. 65:3–10.

Salces-Ortiz J, Vargas-Chavez C, Guio L, Rech GE, González J. 2020. Transposable elements contribute to the genomic response to insecticides in Drosophila melanogaster. Philos. Trans. R. Soc. B 375:20190341–20190341.

Schattner P. 2008. Genomes, Browsers, and Databases: Data-mining Tools for Integrated Genomic Databases. Cambridge University Press

Sienski G, Dönertas D, Brennecke J. 2012. Transcriptional Silencing of Transposons by Piwi and Maelstrom and Its Impact on Chromatin State and Gene Expression. Cell 151:964–980.

Solares EA, Chakraborty M, Miller DE, Kalsow S, Hall K, Perera AG, Emerson JJ, Hawley RS. 2018. Rapid Low-Cost Assembly of the Drosophila melanogaster Reference Genome Using Low-Coverage, Long-Read Sequencing. G3 GenesGenomesGenetics 8:3143–3154.

