## Supplementary Material for "DrosOmics: a comparative genomics browser to explore omics data in natural populations of *D. melanogaster*"

Methods for obtaining and processing the functional -omics data for five *de novo* *D. melanogaster* genomes from European populations.

### Fly stocks and body part dissection

Five *D. melanogaster* strains obtained from the European Drosophila Population Genomics Consortium (DrosEU), were selected according to their different geographical origins: AKA-017 (Akaa, Finland); JUT-011 (Jutland, Denmark); MUN-016 (Munich, Germany); SLA-001 (Slankamen, Serbia) and TOM-007 (Tomelloso, Spain). Flies were reared on fly food medium in a 12:12 h light/dark cycle at 25°C.

Guts, ovaries and heads for each strain were dissected from the same individuals in 1× phosphate buffered saline (PBS) from 4-6 days old females and immediately frozen in liquid nitrogen. 1× Protease inhibitor was added to the 1× PBS for ChIP-seq tissues. Three replicates of 30 females each were processed per tissue and strain.

### RNA extraction, library preparation and sequencing

RNA from guts and ovaries were extracted using GenElute Mammalian genomic total RNA miniprep kit from SIGMA following the manufacturer's instructions.

Head samples were homogenized in 1mL of TRIzol (Invitrogen) using an electric pestle. RNA was purified according to the suggested manufacturer's protocol (Invitrogen).

All samples were DNase-treated using Ambion™ DNase I (#AM2222; Thermo Scientific). RNA was finally precipitated with lithium chloride (LiCl 5M). RNA quality and concentration were assessed by NanoDrop™ (Thermo Fischer) and Bioanalyzer (Agilent). 1.5µg of total RNA from each sample was used for subsequent library preparation and sequencing. Briefly, library preparation was performed using the Truseq Stranded mRNA Sample Prep kit from Illumina following the manufacturer's instructions. Libraries were sequenced using Illumina 125 bp paired-end reads (ranging from 26.4 million reads to 68.8 million reads, Table S2). Sequencing was performed in the CRG Genomics Core Facility in Barcelona.

### ChIP library preparation and sequencing

Guts and ovaries were resuspended in Buffer A1 [60mM KCl, 15mM NaCl, 15mM HEPES pH 7.4, 0.5% Triton ×100, 10mM Sodium Butyrate and 1× Protease Inhibitor Cocktail (1×PI, Roche)] and crosslinked in formaldehyde at a final concentration of 1.8%. They were gently homogenized (∼20 strokes) in a 2mL glass tissue grinder (Dounce B).

Heads were pulverized to a fine powder using an electrical pestle. The fine powder was resuspended in Buffer A1 and crosslinked in formaldehyde at a final concentration of 1.8%. In all cases, after 10 minutes rotating in a wheel at room temperature, crosslinking was stopped by the addition of Glycine at a final concentration of 0.25M.

All the following steps were performed at 4°C: samples were centrifuged at 4000g for 5 minutes. The pellet was subsequently washed with Buffer A1 (3 times) and Lysis buffer without SDS [140mM NaCl, 15mM HEPES pH 7.4, 1mM EDTA, 0.5mM EGTA, 1% Triton ×100, 0.1 Sodium deoxycholate, 10mM Sodium butyrate and 1× Protease Inhibitor Cocktail (1×PI, Roche)]. Cells were then incubated for 2 hours with 0.2 mL of lysis buffer (HEPES 15mM, EDTA 1mM, EGTA 0.5 mM, Sodium Butyrate 10mM, SDS 0.5%, Sodium deoxycholate 0.1%, N-Lauroylsarcosine 0.5%, Triton ×100 1%, and NaCl 140mM).

Chromatin was sheared with a Bioruptor sonicator water bath (Diagenode, Liège, Belgium). All samples were sonicated with cycles of 30-s on/30-s off at high power. To generate random fragments from 250 to 500 bp ovaries were sonicated for 15 cycles, guts for 20 cycles and heads for 35 cycles and fragments sizes were checked with bioanalyzer (Agilent). Chromatin was resuspended by centrifugation at 10,000g for 10 minutes at 4ºC.

All the subsequent steps were performed using the Magna ChIP G chromatin immunoprecipitation kit (Millipore). All the buffers and reagents used were provided by the kit and the manufacturer's instructions were followed. We separated 20µl of the chromatin as input and stored it at 4°C. The remaining 180µl were divided in 2 aliquots (each one for each antibody used). To each aliquot we added 3µg of antibody [H3K9me3 (Abcam #ab8898) and H3K27ac (Abcam #ab4729)] plus 20µl of magnetic beads and dilution buffer up to 530µl. We incubated the samples overnight at 4°C in an agitation wheel. After incubation, beads were sequentially washed with Low salt buffer, High salt buffer, LiCl complex buffer, and TE buffer provided by the kit. Chromatin from the immunoprecipitations (IPs) were eluted in 0.5mL of elution buffer. Proteinase K was added both for IP and inputs and incubated at 65°C in a shaker at 300 rpm overnight. After incubation, chromatin was purified by phenol:chloroform using MaXtract High Density tubes (Qiagen). Samples were eluted in 20µl of DEPEC water and stored at -20ºC until further processed.

Libraries were performed using TruSeq ChIP Library Preparation Kit following manufacturer’s instructions. Sequencing was carried out in an Illumina Hiseq 2500 platform, generating 50 bp single-end reads (ranging from 22.2 million reads to 59.1 million reads, Table S2).

**Data availability**

RNA-seq and ChIP-seq raw data is available from the NCBI Sequence Read Archive (SRA) database under BioProject PRJNA643665.

### Bioinformatic processing

The following protocols for processing RNA-seq and ChIP-seq data were applied to all samples compiled in DrosOmics.

#### RNA-seq alignment and isoform quantification

We align each RNA-seq to their corresponding reference genome following the protocol described in Pertea et al. (2016), which uses HISAT2 (v2.2.1; Kim et al. 2019). To improve the accuracy of the mapping, we used the gene annotations of each genome. For that, we used extract_splice_sites.py and extract_exons.py python scripts included in the HISAT2 package, to extract splice site and exon information from the gene annotation file. Next, we build the HISAT2 index using hisat2-build (argument -p 12) providing the splice sites and exons in the -ss and -exon arguments. Next, we performed the mapping of the RNA-seq reads with HISAT2. The output sam files were sorted and transformed into bam files using SAMtools (v1.6; Li et al. 2009). Next, we used StringTie (v2.1.2; Pertea et al. 2015) for the assembly of isoforms using the gene annotations of each strain. To obtain the gene count estimations normalized by the trimmed mean of M-values method (TMM; Robinson et al. 2010), for better comparison between samples, we first used the prepDE.py script from stringtie to obtain a matrix of gene counts, changing the length parameter -l 125, as our Illumina reads are of 125 bp. Then, we used the edgeR R package (v3.28.0, Robinson et al. 2010) to first create a DGEList object from the table of counts using the DGEList() function. Next, we used calcNormFactors() function using the TMM method to calculate the normalization factors to scale the raw library sizes. Finally, we used the function cpm() to obtain the normalized counts using the TMM normalization factors.

#### Z-score estimation for *Myo31DF* gene in TOM-007 and JUT-011 genomes

We calculate the expression z-score as: mean of gene expression of gene with TE (TOM-007, considering the three biological replicates) minus the mean expression of gene without the TE (JUT-011), divided by the standard deviation of both group of genes genes (Equation 1). A negative z-score indicates that the gene associated with the TE insertion (TOM-007) has a lower expression compared to the gene in the genome without the insertion (JUT-011). The p-value is calculated using the cumulative distribution function cbd() from the distributions3 package in R (v0.1.2; Hayes et al. 2022) from a normal distribution.

$z=\frac{\underline{x_{1}}-\underline{x_{2}}}{\sqrt{\frac{\sigma_{1}^{2}}{n_{1}}+\frac{\sigma_{2}^{2}}{n_{2}}}}$	Equation 1

#### RNA-seq coverage track files

To create RNA-seq coverage profile files for visualization in the browser, we used bamCoverage (v3.5.1) from the deepTools package (Ramírez et al. 2016). We transformed the bam files into bigwig files with options --binSize 10 to set the size of the bins in bases and --normalizeUsing BPM to normalize it by bins per million mapped reads (BPM).

#### ChIP-seq processing: enrichment signals and peak calling

ChIP-seq reads were processed using fastp (v0.20.1; Chen et al. 2018) to remove adaptors and low-quality sequences. Processed reads were mapped to the corresponding reference genome using the readAllocate function (with option chipThres = 500) of the Perm-seq R package (v0.3.0; Zeng et al. 2015) with bowtie (v1.2.2; Langmead et al. 2009) as the aligner and the CSEM program (v2.3; Chung et al. 2011) in order to try to define a single location to multi-mapping reads. In all cases, bowtie was used with default parameters selected by Perm-seq. Then we used the ENCODE ChIP-Seq caper pipeline (v2; Landt et al. 2012) in histone mode, using bowtie2 as the aligner, disabling pseudo replicate generation and all related analyses (chip.true_rep_only=TRUE) and pooling controls (chip.always_use_pooled_ctl=TRUE). The MACS2 peak caller was used with default settings (Gaspar 2018). We used two outputs generated by MACS2: the narrowPeak files with the bigWig files with the signal value tracks (*call-macs2_signal_track* file) and the called peaks (*call-peak* file).

#### ChIP-seq track files

We used bigWigToBedGraph to transform the signal value tracks into bed files (Kent et al. 2010). Next, we used bedtools makewindows (v2.30; Quinlan and Hall 2010) to create bins of 10 bases (-w 10). Next, bedmap from the BEDOPS package (v2.4.39; Neph et al. 2012) was used to create a bed file with the mean signal value of each bin (--echo --mean options). Finally, bedGraphToBigWig was used to transform the bed file to bigWig. We did that to keep both RNA-seq and ChIP-seq tracks in 10-bp bins.

Finally, to visualize the peaks, we used bedToBigBed with options -type=bed4+6 -as=narrowPeak.as to create a bed file with the bigNarrowPeak information available in <http://genome.ucsc.edu/goldenPath/help/examples/bigNarrowPeak.as>.

### Additional references

Chen S, Zhou Y, Chen Y, Gu J. 2018. fastp: an ultra-fast all-in-one FASTQ preprocessor. *Bioinformatics* 34:i884–i890.

Chung D, Kuan PF, Li B, Sanalkumar R, Liang K, Bresnick EH, Dewey C, Keleş S. 2011. Discovering Transcription Factor Binding Sites in Highly Repetitive Regions of Genomes with Multi-Read Analysis of ChIP-Seq Data. *PLOS Comput. Biol.* 7:e1002111.

Gaspar JM. 2018. Improved peak-calling with MACS2. *bioRxiv*:496521v1.

Hayes A, Moller-Trane R, Jordan D, Northrop P, Lang MN, Zeileis A. 2022. distributions3: Probability Distributions as S3 Objects.

Kent WJ, Zweig AS, Barber G, Hinrichs AS, Karolchik D. 2010. BigWig and BigBed: enabling browsing of large distributed datasets. *Bioinforma. Oxf. Engl.* 26:2204–2207.

Kim D, Paggi JM, Park C, Bennett C, Salzberg SL. 2019. Graph-based genome alignment and genotyping with HISAT2 and HISAT-genotype. *Nat. Biotechnol.* 37:907–915.

Landt SG, Marinov GK, Kundaje A, Kheradpour P, Pauli F, Batzoglou S, Bernstein BE, Bickel P, Brown JB, Cayting P, et al. 2012. ChIP-seq guidelines and practices of the ENCODE and modENCODE consortia. *Genome Res.* 22:1813–1831.

Langmead B, Trapnell C, Pop M, Salzberg SL. 2009. Ultrafast and memory-efficient alignment of short DNA sequences to the human genome. *Genome Biol.* 10:R25.

Li H, Handsaker B, Wysoker A, Fennell T, Ruan J, Homer N, Marth G, Abecasis G, Durbin R, 1000 Genome Project Data Processing Subgroup. 2009. The Sequence Alignment/Map format and SAMtools. *Bioinformatics* 25:2078–2079.

Neph S, Kuehn MS, Reynolds AP, Haugen E, Thurman RE, Johnson AK, Rynes E, Maurano MT, Vierstra J, Thomas S, et al. 2012. BEDOPS: high-performance genomic feature operations. *Bioinformatics* 28:1919–1920.

Pertea M, Kim D, Pertea G, Leek JT, Salzberg SL. 2016. Transcript-level expression analysis of RNA-seq experiments with HISAT, StringTie, and Ballgown. *Nat. Protoc.* 11:1650–1667.

Pertea M, Pertea GM, Antonescu CM, Chang T-C, Mendell JT, Salzberg SL. 2015. StringTie enables improved reconstruction of a transcriptome from RNA-seq reads. *Nat. Biotechnol.* 33:290–295.

Quinlan AR, Hall IM. 2010. BEDTools: a flexible suite of utilities for comparing genomic features. *Bioinformatics* 26:841–842.

Ramírez F, Ryan DP, Grüning B, Bhardwaj V, Kilpert F, Richter AS, Heyne S, Dündar F, Manke T. 2016. deepTools2: a next generation web server for deep-sequencing data analysis. *Nucleic Acids Res.* 44:W160-165.

Robinson MD, McCarthy DJ, Smyth GK. 2010. edgeR: a Bioconductor package for differential expression analysis of digital gene expression data. *Bioinformatics* 26:139–140.

Zeng X, Li B, Welch R, Rojo C, Zheng Y, Dewey CN, Keleş S. 2015. Perm-seq: Mapping Protein-DNA Interactions in Segmental Duplication and Highly Repetitive Regions of Genomes with Prior-Enhanced Read Mapping. *PLOS Comput. Biol.* 11:e1004491.
